# Biofabrication of nanocomposite-based scaffolds containing human bone extracellular matrix for the differentiation of skeletal stem and progenitor cells

**DOI:** 10.1101/2023.04.07.536074

**Authors:** Yang-Hee Kim, Janos M Kanczler, Stuart Lanham, Andrew Rawlings, Marta Roldo, Gianluca Tozzi, Jonathan I. Dawson, Gianluca Cidonio, Richard O.C Oreffo

**Affiliations:** Bone and Joint Research Group, Centre for Human Development, Stem Cells and Regeneration, Institute of Developmental Sciences, Faculty of Medicine, University of Southampton, Southampton, SO16 6YD, United Kingdom; 3D Microfluidic Biofabrication lab, Center for Life Nano- & Neuro-Science (CLN2S), Fondazione Istituto Italiano di Tecnologia, Viale Regina Elena 291, 00161, Rome, Italy; School of Pharmacy and Biomedical Science, St Michael’s Building, White Swan Road, University of Portsmouth, PO1 2DT Portsmouth, United Kingdom; School of Engineering, Faculty of Engineering & Science, Chatham Maritime, University of Greenwich, ME4 4TB Greenwich, United Kingdom

**Keywords:** nanoclay, bone, 3D bioprinting, extracellular matrix

## Abstract

Autograft or metal implants are routinely used in skeletal repair but can fail to provide a long-term clinical resolution, emphasising the need for a functional biomimetic tissue engineering alternative. An attractive sustainable opportunity for tissue regeneration would be the application of human bone waste tissue for the synthesis of a material ink for 3D bioprinting of skeletal tissue.

The use of human bone extracellular matrix (bone-ECM) offers an exciting potential for the development of an appropriate micro-environment for human bone marrow stromal cells (HBMSCs) to proliferate and differentiate along the osteogenic lineage. Extrusion-based deposition was mediated by the blending of human bone-ECM (B) with nanoclay (L, Laponite^®^) and alginate (A) polymer, to engineer a novel material ink (LAB). The inclusion of nanofiller and polymeric material increased the rheological, printability, and drug retention properties and, critically, the preservation of HBMSCs viability upon printing. The composite human bone-ECM-based 3D constructs containing vascular endothelial growth factor (VEGF) enhanced vascularisation following implantation in an *ex vivo* chick chorioallantoic membrane (CAM) model. Addition of bone morphogenetic protein-2 (BMP-2) with HBMSCs further enhanced vascularisation together with mineralisation after only 7 days.

The current study demonstrates the synergistic combination of nanoclay with biomimetic materials, (alginate and bone-ECM) to support the formation of osteogenic tissue both *in vitro* and *ex vivo* and offers a promising novel 3D bioprinting approach to personalised skeletal tissue repair.

**Graphical Abstract:** Engineering nanoclay-based bone ECM novel bioink for bone regeneration. Human bone trabecular tissue was demineralised, decellularised and blended with nanoclay (Laponite®) and alginate after digestion. The resulting ink was investigated for printability following rheological and filament fusion investigation. The microstructural arrangement of the blends was examined together with viability and functionality of bioprinted HBMSCs. Finally, the ability of the novel blend to support drug release ex vivo in a CAM model was determined confirming the potential of the bone ECM ink to support bone formation.

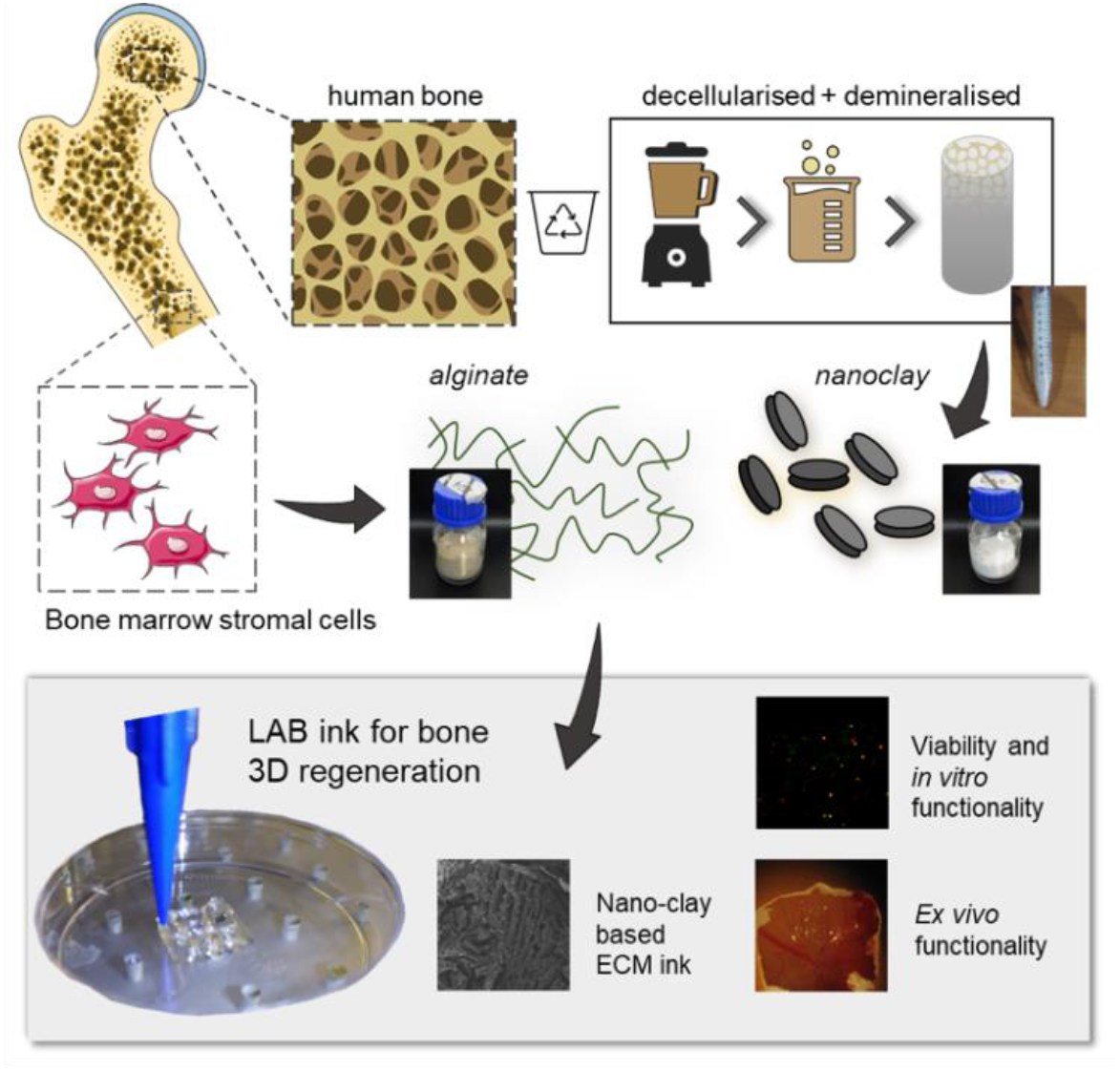

## 1. Introduction

Skeletal tissue engineering (TE) aims to provide functional tools for the repair of damaged or diseased bone tissue. Over the last decade, biofabrication approaches for TE have explored a number of biomaterials able to degrade over time, support cell delivery and sustain the release of biological agents of interest for the repair ^1–3^ or the modelling ^4^ of bone.

However, material inks formulated using either natural or synthetic platforms have, to date, proved unable to fully support skeletal repair failing to resemble/recapitulate the native bone micro-environment ^5–8^. Recently, organic nano-fillers have offered significant promise to enhance printability and skeletal functionality^3^. Particularly, nanoclays have been employed to engineer a library of material inks capable of sustaining skeletal stem and progenitor cell viability and differentiation *in vitro*^9^, *ex vivo*^10,11^ and *in vivo*^12^. These nanoclay composites provide a powerful tool for the engineering of a rapidly evolving skeletal micro-environment. However, nanocomposite materials are unable to fully recapitulate and mimic the native skeletal micro-environment, limiting the biomimetic platform for stem-progenitor cell differentiation and skeletal maturation.

The native bone tissue physico-chemical composition remains the ideal material for skeletal repair. Indeed, bone extracellular matrix (ECM) contains a plethora of growth factors (GFs) (e.g. vascular endothelial growth factor (VEGF), bone morphogenetic protein-2 (BMP-2) and others) as well as polymeric constituents (e.g. collagen), essential for the development and repair of skeletal tissue ^13,14^. For a number of years, autologous and allogenic bone grafts have been routinely used clinically to repair large skeletal defects. Impaction bone graft is used to repair segmental defects harnessing cadaveric tissue. Nevertheless, (i) the scarcity of available bone tissue, (ii) the lack of donor-to-donor compatibility and, (iii) the functional ability to match defect architecture and regenerative capacity have limited the use of human-derived bone grafts. Moreover, the inability of impaction bone grafts to fully facilitate bone regeneration remains a limitation. A potential solution to these issues is the application of biomaterials engineered from native skeletal tissue. The use of decellularised allografts would allow the isolation of native ECM material together with the removal of anyallogenic cellular components and epitopes that could trigger an immune response upon implantation. Recent advancements in decellularization techniques have facilitatedthe preparation of ECM derived from tissue previously difficult to digest and process. However, human-based decellularised ECM tissues have, to date, not been successfully applied in skeletal tissue engineering applications., xenogenic ECM materials have been explored as printable inks to support tissue-specific repair harnessing physiological mechanisms from naturally-derived matrices^15^. A number of studies in the last decade^16^ have explored the possibility to isolate ECM-based materials from animal tissues including cardiac^17^ and liver^18^. Nevertheless, animal-derived ECM material inks remain a significant challenge for human application given species differences and immunologic considerations. Tissue-derived ECM materials have failed to function effectively as reproducible bioprintable platform due to: (i) complex matrix derivation steps typically required, involving acidic components and extensive filtering procedures, (ii) the poor viscoelastic properties of the derived materials limiting extrusion-based bioprinting approaches and, (iii) species differences between ECM composition of animal and human sources and the accompanying host-immune response issues encountered upon implantation ^19,20^. Thus, a human-sourced ECM material ink could provide a step-change in the bioink design paradigm, offering an innovative approach to personalised skeletal regenerative medicine ^21^.

The current study demonstrates the printability, *in vitro* stability and *ex vivo* functionality of a novel human bone ECM-based bioink composite. The inclusion of a nanoclay filler was found to improve the physico-chemical properties limiting the swelling rate and porosity while enhancing the material viscosity profile (**Figure 1a**). 3D printing of the HBMSC-laden bone-ECM material resulted in a stable culture construct that supported cell growth and promoted skeletal cell functionality *in vitro* (**Figure 1b**) and *ex vivo* (**Figure 1c**). The inclusion of nanoclay particles was supportive for *ex vivo* drug retention compared to the clay-free controls, providing a platform able to support vascular and bone regeneration. This biomimetic nanocomposite material offers a promising 3D bioprinting approach for personalised skeletal tissue repair.

**Figure 1.**
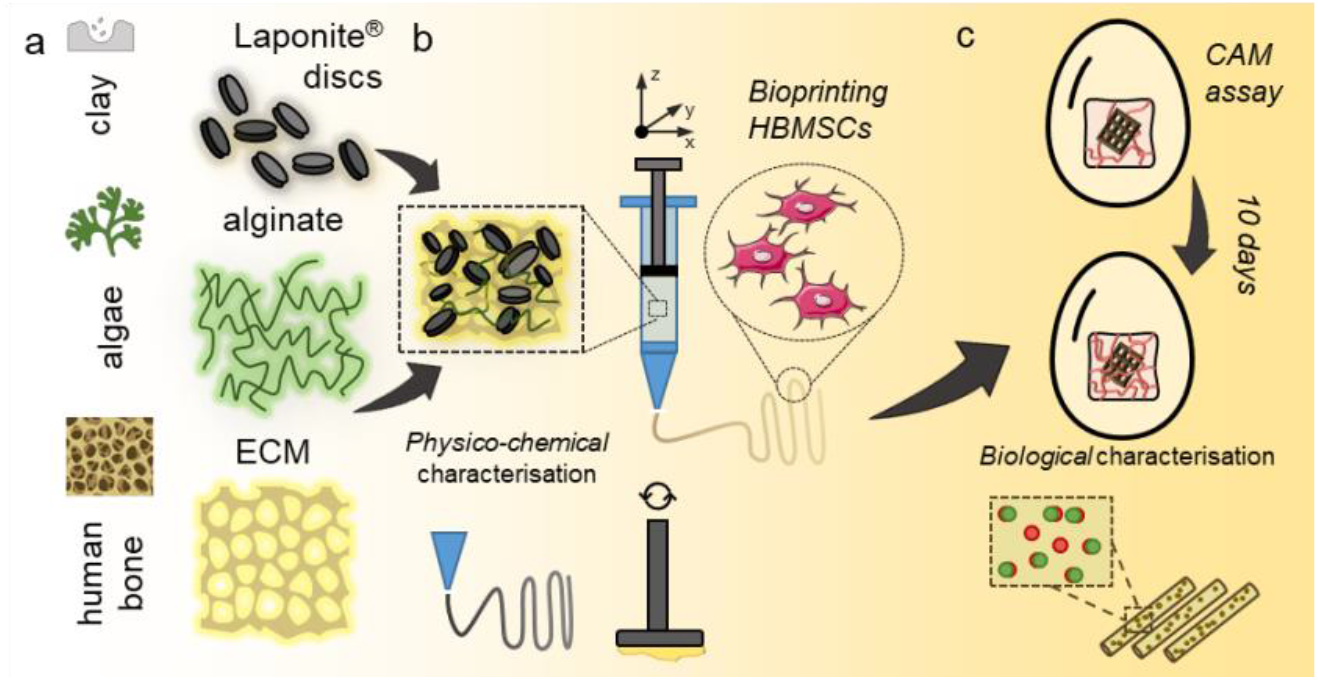
A novel biomaterial ink system engineered from the combination of nanoclay discs, alginate and a novel human bone demineralised and decellularised ECM. The nanocomposite ink rheological properties were investigated, along with the ability of the nanocomposite ink to be printed with elevated resolution in three dimensions. The inclusion of HBMSCs allowed analysis of viability and differentiation over 21 days, as well as evaluation and demonstration of3D functionality in a CAM model.

## 2. Materials and Methods

### 2.1 Nanocomposite hydrogels preparation

Nanocomposite hydrogels were prepared in a sterile Class II cell culture hood. Laponite^®^ (L, XLG grade, BYK Additives & Instruments, UK) was allowed to disperse at either 3 or 4 wt% (30 or 40 mg ml^−1^ respectively) in suspension in deionised water (DW) for 3 h under constant stirring until clear, followed by UV sterilisation. Bone ECM was prepared following a previously employed protocol ^21^. Briefly, cancellous bone fragments were collected from donated femoral heads from patients undergoing total hip-replacement with full national ethical approval following informed patient consent (Southampton General Hospital, University of Southampton under approval of the Southampton and Southwest Hampshire Research Ethics Committee (Ref No. 194/99/1)), using a bone nipper and washed with 2% penicillin/streptomycin (P/S). Bone fragments were ground to a fine powder and stirred in 0.5 N HCl at room temperature for 24 h to allow complete demineralisation as previously reported ^22^. The demineralised bone matrix (DBM) was fractionated using a 45 µm-pore sieve and washed with deionized water (DW). A 1:1 mixture of chloroform and methanol was used to treat the DBM for 1h and to facilitate extraction of the lipidic portion. The lipid-free DBM was subsequently lyophilised overnight and stored at -20 °C for future use. To deplete the cellular component of the DBM, a 0.05% Trypsin and 0.02% EDTA solution was added to the DBM and left to stir at 37 °C in a 5% CO_2_ incubator for 24 h. The decellularised DBM was further rinsed and treated in pepsin solution (20 mg ECM/1 ml of pepsin solution) under constant agitation at room temperature for 7 days, followed by centrifugation. The supernatant (referred to as decellularised matrix – ECM (B)) was collected and lyophilised.

The lyophilised bone ECM (B) was added at a concentration of 10 mg ml^−1^ to a Laponite^®^ (L) suspension. Following 2 h stirring at room temperature, alginate (A, alginic acid sodium salt from brown algae, 71238, Sigma, UK) was added to L-B suspension and homogenised with a spatula for 8-10 min to allow alginate inclusion. The combinations of LAP, alginate and bone-ECM examined in this study are detailed in Table 1. LAB ink was stored at room temperature and printed the following day.

**Table 1.**
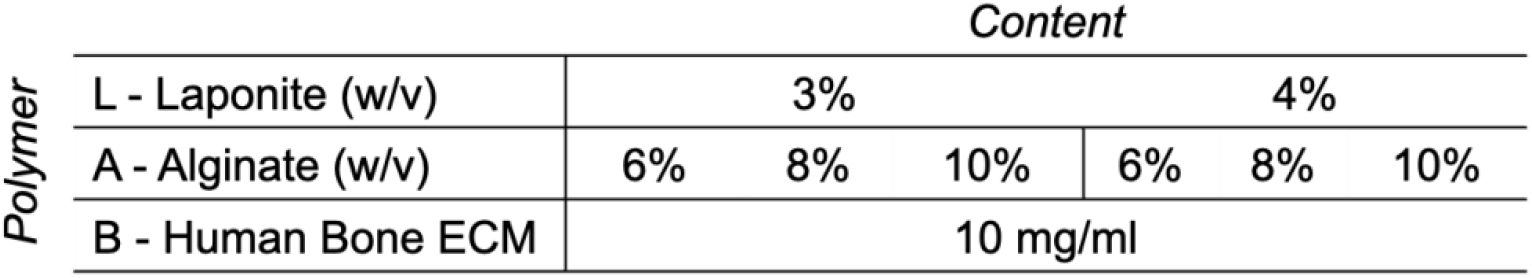
Schematic of composite ink combinations used in this study. Laponite (L) and Alginate (A) were mixed with human bone ECM (B) to generate material composites for further physico-chemical characterisation.

### 2.2 Physico-chemical characterisation

#### 2.2.1 Mass loss and swelling studies of nanocomposite hydrogels

To investigate the effect of Laponite^®^ (L) on alginate (A) and bone ECM (B), the mass loss and swelling ratio of nanocomposite gels was investigated as previously detailed ^23^.

LAB hydrogels with various concentrations of L and A (Table 1) were prepared and cast in 500 µl moulds. To obtain the initial wet mass, samples (n=3) were weighed before (*m*_*initial*_) and after cross-linking (*m*_*initial,t*0_). LAB samples (n=3) were lyophilized to obtain their dry weights (*m*_*dry,t*0_). The macromer fraction was calculated as follows: (1)

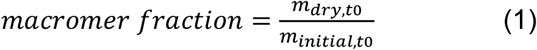

The remaining samples (n=3) were incubated at 37 °C in phosphate buffered saline (PBS, Thermo-Fisher) or Hank’s balanced salt solution (HBSS, Thermo-Fisher). The samples were reweighed (*m*_*swollen*_) after 24 h. LAB samples were subsequently lyophilized and weighed (*m*_*dry*_). The sol fraction was calculated using equation (2) and (3). Mass swelling ratio (q) was then calculated using equation (4).

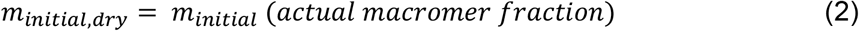

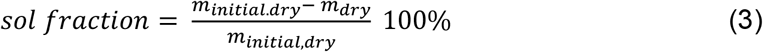

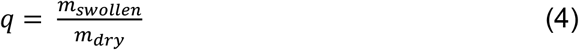

#### 2.2.2 Scanning electron microscopy

Scanning electron microscopy (SEM, FEI Quanta 200 FEG) at a voltage of 5 kV (spot size 3) was used to image the acellular gels. Samples were dehydrated using a freeze-drier (Lablyo Mini, Froze in Time Ltd., UK) for 12 h and platinum-coated to allow SEM analysis (Q150TES, sputter coater, UK). Porosity was calculated from SEM images (n=3) using Image J analysis 24.

#### 2.2.3 Rheological measurements of nanocomposite hydrogel properties

Rheological measurements of nanocomposite hydrogels were carried out using a cone-plate rheometer (Anton Parr, MCR92) at room temperature with a 0.1 mm gap. A range of shear rate between 1 and 100 s^−1^ with a linear increase was applied to measure viscosity (Pa s) of the LAB ink formulations and controls. The stable value for the viscosity of the inks was measured applying a constant shear rate (10 s^−1^) for 720 s. Considering a viscoelastic behaviour at 1% shear strain, frequency sweeps were carried out over a range between 0.01 and 100 s^−1^. Storage and Loss moduli of controls and LAB material inks were acquired at 1% shear strain.

### 2.3 Printing fidelity

The fidelity of filament deposition was assessed as previously published ^25^. Briefly, three layers were deposited consequently layering strands at an increasingly larger distance. Scaffolds (n=3) were imaged immediately following deposition using a light stereomicroscope (Zeiss) equipped with Canon Powershot G2 camera. Acquired images were analysed using Image J, identifying fused segment length (*fs*), filament thickness (*ft*) and filament distance (*fd*). Results were plotted as the ratio of *fs* and *ft* as a function of the *fd*.

### 2.4 Printing of nanocomposite ink

LAB inks were deposited to investigate printing fidelity as previously published ^25^. Briefly, LAB inks were printed in a winding pattern with an exponentially increasing strand distances and imaged (Stemi DV4, Zeiss, UK) immediately after printing. Images were analysed with Image J software obtaining actual strand distance, fused segment length and strand width. The filament fusion test was then graphed based on the quotient of segment length and strand width as a function of strand distance. An in-house built bioprinter ^11^ was used to deposit acellular and cell-laden LAB inks, using a 410 µm nozzle (Fisnar Europe, UK). Multi-layers scaffolds of 10 × 10 mm^2^ with an alternating pattern (ABAB, 0°/90°), a layer height of 350 μm and a strand distance of 2 mm were printed. A solution of 100 mM CaCl_2_ was used to crosslink the printed structures for 10 min. Scaffolds for viability (total n=12) and functionality (total n=16) investigations were printed with n=3 scaffolds used at each time point.

### 2.5 Cell isolation, encapsulation and printing

Unselected HBMSCs were isolated as previously described [12] from patients undergoing total hip-replacement with full national ethical approval following informed patient consent (Southampton General Hospital, University of Southampton under approval of the Southampton and Southwest Hampshire Research Ethics Committee (Ref No. 194/99/1)). Briefly, to remove excessive fat, the bone marrow aspirate was resuspended and washed in alpha modified eagle’s medium (α-MEM), filtered through a 40-µm cell strainer and layered on LymphoPrep™ (Lonza) with a density centrifugation at 2200 RPM (800 G) for 40 min at 18°C. The portion of bone marrow mononuclear cells (BMMNCs) was isolated and plated in cell culture flasks and maintained at 37 °C and 5 % CO_2_ balanced air with α-MEM supplemented with 10% v/v fetal bovine serum (FBS), 100 U ml^−1^ penicillin and 100 µg ml^−1^ streptomycin (Pen/Strep). Cells were passaged at approximately 80 % cell confluency using collagenase IV (200 mg ml^−1^) in serum-free media followed by Trypsin-ethylenediaminetetra-acetic acid (TE) solution treatment. HBMSCs were used for experimental studies at passage two. To visualise cells after printing for viability studies, cells were suspended at a density of 1 × 10^6^ cells ml^−1^ in serum-free culture medium and labelled with Vybrant^®^ DiD (Cell-Labeling Solution, V22887, Molecular Probes) following manufacturer protocol. In brief, the cell suspension supplemented with DiD was incubated for 20 min at 37 °C. Following centrifugation, the supernatant was removed, and the stained cell pellet washed in serum-free culture media. Cells were suspended in 50 µl of serum-free media and added to the material ink. The bioink was mixed with a sterile spatula prior to loading the syringe for printing. Cell printing was carried out using a 410 µm nozzle (Fisnar Europe, UK) fabricating 10 × 10 mm^2^ scaffolds with an alternating layer pattern (ABAB, 0°/90°). Following deposition, 3D printed scaffolds were incubated for 10 min in sterile 100 mM CaCl_2_ solution and then incubated at 37 °C and 5 % CO_2_ balanced air. Cell-laden scaffolds for viability and functionality were printed in triplicates for each time point using DiD-stained and unstained bioinks, respectively.

### 2.6 Viability and functionality analysis

Cell viability was investigated after 1, 7 and 21 days of culture using confocal imaging as previously described ^9^. In brief, samples were washed twice with 1× HBSS. Scaffolds were then incubated in a diluted serum-free culture media solution of Calcein AM (C3099, Invitrogen, Thermo Fisher Scientific, UK) at 37 °C and 5 % CO_2_ balanced air for 1 h, following the manufacturer’s protocol. Living cells were stained by both Calcein AM and DiD (green and red). Cells non-metabolically active (dead) were stained by DiD in red. Scaffolds were imaged using a confocal scanning microscope (Leica TCS SP5, Leica Microsystems, Wetzlar, Germany). Image analysis was carried out using Image J. Cell density was calculated normalising the number of viable cells with the volume of interest (VOI).

Cell-laden scaffolds were cultured in basal (α-MEM supplemented with 10 % (v/v) FBS and 1% Pen/Strep) and osteogenic (α-MEM supplemented with 10 % (v/v) FBS and 1% Pen/Strep, 100 µM ascorbate-2-phosphate (AA2P, Sigma-Aldrich), 10 nM dexamethasone (Dex, Sigma-Aldrich) and 10 nM vitamin D (1α,25-OH2-Vit D3, Sigma-Aldrich)) conditioned media.

Alkaline phosphatase (ALP) staining was carried out after 1, 7, 21 days of culture at 37 °C in 5 % CO_2_ balanced air. Samples were washed twice with 1× HBSS and fixed in 95% ethanol for 10 min. Scaffolds were left to dry while ALP staining solution was prepared combining Naphthol (AS-MX Phosphate Alkaline Solution, 85-5, Sigma-Aldrich, UK) and Fast Violet Salt (F1631 Sigma-Aldrich, UK) solubilised in DW. Samples were incubated at 37 °C for 1 h with the ALP staining solution and the reaction was stopped by dilution of ALP solution with HBSS. Stained scaffolds were stored at 4 °C overnight and imaged the following day using a Zeiss Axiovert 200 (Carl Zeiss, Germany).

### 2.7 Modelling absorption and release

Protein absorption and release study was carried out as previously reported ^10^. Model proteins lysozyme from chicken egg (Sigma-Aldrich, UK) and bovine serum albumin (BSA, Sigma-Aldrich, UK) were solubilised in Hank’s Balanced salt solution (HBSS, Thermo-Fisher, UK) at 10 µg·ml^−1^ and 100 µg·ml^−1^, respectively. To investigate the influence of nanoclay particles on drug release, nanocomposite (LAB) and Laponite-free controls (AB) were used to print 3D scaffolds to allow absorption of compounds of interest following ionic crosslinking. The 3D printed constructs (n=3) were soaked in lysozyme or BSA for 1 h, then release monitored over 24h. BSA and lysozyme were quantified with a quantification kit (Rapid-kit, Sigma-Aldrich, UK) using GloMax Discover microplate reader (Promega). The supernatant was collected after 1, 2, 4, 8, 10, 20 and 24 hours following adsorption. Collagenase D (from Clostridium histolyticum, Roche Diagnostics GmbH) was added following 24 h from adsorption, to stimulate material degradation and cargo release. BSA and lysozyme release were quantified after 1, 2, 4, 8, 10, 20 and 24 hours.

### 2.8 Chick chorioallantoic membrane (CAM) model

#### *Scaffold fabrication for ex vivo* vascularisation

Scaffolds with nanoclay and LAP-free were 3D printed, crosslinked following 10 min exposure to 100 mM CaCl_2_ and allowed to adsorb for 30 min with recombinant human vascular endothelial growth factor (rhVEGF 165, PeproTech, USA) at 100 μg mL^−1^ at 4 °C. 3D printed constructs were washed three times with HBSS 1x prior storage overnight at 4 °C.

#### *Scaffold fabrication for ex vivo cell delivery and* mineralisation

Nanoclay-based and LAP-free 3D scaffolds were fabricated and implanted immediately after adsorption of recombinant human bone morphogenetic protein-2 (rhBMP-2) at 10 µg ml^*-*^1 for 30 min at 4 °C. Scaffolds were washed in HBSS 1X for three times before implantation.

#### CAM implantation, extraction and Chalkley score

The CAM *ex vivo* model was used to evaluate vascularisation and mineralisation. Animal studies were conducted in accordance with Animals Act 1986 (UK), under Home Office Approval UK (PPL P3E01C456). Fertilised eggs were maintained in a rotating Hatchmaster incubator (Brinsea, UK) for 10 days at 37 °C and 60% humidity. 3D printed scaffolds were implanted at day 10 post-fertilisation. The implantation was carried out under a class II laminar flow hood operating a 2 cm^2^ window on the eggshell. Constructs were laid on the CAM membrane and eggshell windows sealed with sterile parafilm. Chicken eggs were then incubated in a non-rotating incubator for 7 days. Developing chick embryos were inspected daily via candling monitoring growth and viability. Following 7 days of incubation, samples were harvested, and CAM integration assessed using a stereomicroscope equipped with a digital camera (Canon Powershot G2). The overlap morphometry analysis was performed on extracted samples as previously described^10^. Briefly, implanted samples were screened for vascular penetration by superimposing the Chalkley graticule and the afferent integrated CAM vasculature. The numbers of counted vessels colliding with the points on the graticule were assessed blind, and each sample assessed/counted 3 times. Samples were collected and fixed in 4% paraformaldehyde (PFA) overnight, before further processing for histological analysis. Afferent vessels diameter was evaluated following processing of stereomicroscope images using Image J software analysis.

#### Mineral deposit formation

The deposition of mineral tissue was assessed using micro-computed tomography (micro-CT, Bruker Skyscan 1176). Scans were performed on samples fixed with 4% PFA following further wash in HBSS before imaging. A pixel size of 35 µm, 65 kV, 385 µA, 0.7° rotation step, 135 ms exposure, and an aluminium filter (Al) of 1 mm were used. CT reconstructions were obtained via NRecon (Bruker), quantitative analysis was performed using CTAn software (Bruker) to assess the average mineral density. Bone phantoms with pre-determined bone density (0.25 g cm^-3^ and 0.75 g cm^-3^) were used as reference and CT scan calibration.

### 2.9 Histological analysis

Samples explanted from *ex vivo* CAM assay were fixed in 4% PFA solution overnight at 4°C then embedded in paraffin for further processing. A microtome was used to produce tissue sections of 8 μm. Goldner’s Trichrome, Alcian Blue & Sirius Red and Von Kossa staining were carried out following previously employed protocols ^26^. Slides were imaged the following day using a Zeiss Axiovert 200 (Carl Zeiss, Germany).

#### 2.10 Statistical analysis

Experimental studies were evaluated by one-way and two-way ANOVA using Bonferroni’s multiple comparison tests. Analysis was carried out with GraphPad Prism 9.0 and significance set at p<0.05.

## 3. Results

### 3.1 Physico-chemical and mechanical properties of nanoclay-based hydrogels

The physico-chemical properties of bone ECM nanocomposite inks were investigated following printing and maintenance in PBS and HBSS buffers. A range of material composites was explored by varying the LAP concentration from 3% to 4% wt, and the alginic acid inclusion between 6% and 10% wt. The concentration of bone ECM was kept constant as the percentage of inclusion (10 mg/ml) was fixed. The sol fraction (**Figure 2a**) decreased with the increase in alginate concentration, in PBS (**Figure 2a-i**) and HBSS (**Figure 2a-ii**) with a significant reduction between L3A6B and L3A10B. This was consistent with results obtained for alginate controls both in sol fraction (**Supplementary Figure 1a**) and mass swelling ratio data generated (**Supplementary Figure 1b**).

**Figure 2.**
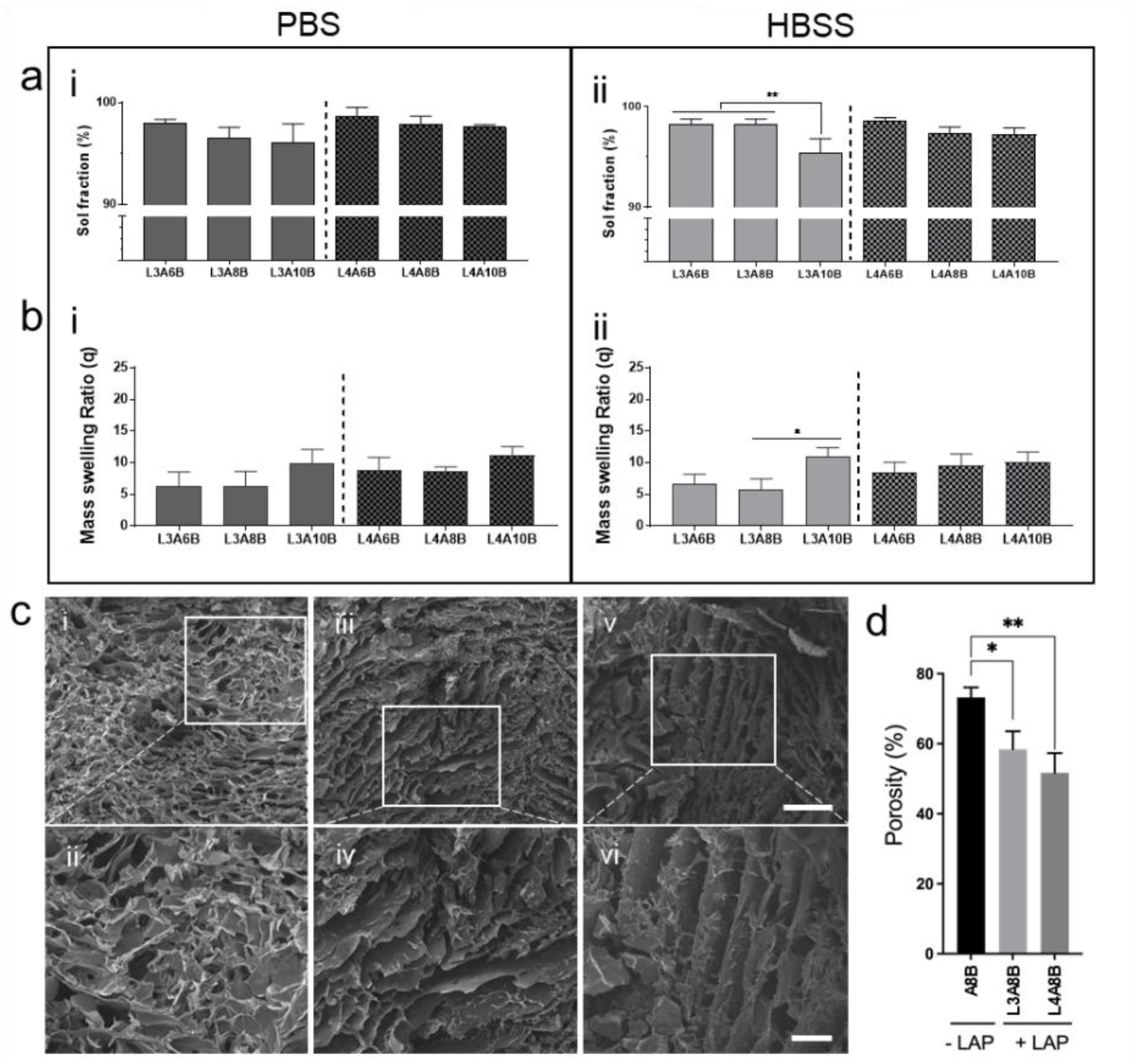
Physical characterisation of composite inks. (a) Sol fraction and (b) mass swelling ratio analysis of scaffolds both in (a-i, b-i) PBS and (a-ii, b-ii) HBSS. SEM micrographs of A8B (c-i,ii) and L4A8B (c-iii,iv) scaffolds. Pore analysis (d) of A8B, L3A8B and L4A8B via Image J measurements. Scale bars: (c-i,iii) 500 µm (c-ii,iv) 200 µm. Statistical significance assessed by unpaired t-test (Welch-corrected). Mean ± S.D. n=3, *p<0.05, **p<0.01

The mass swelling ratio (q) revealed a non-significant increase in swelling as the alginate fraction was increased in the nanocomposite ink in both PBS (**Figure 2b-i**) and HBSS (**Figure 2b-ii**). Controls in PBS (**Supplementary Figure 1c**) and HBSS **(Supplementary Figure 1d**) showed a significant decrease in sol fraction and a proportional increase in swelling ratio as LAP content increased. The microstructural arrangement of Laponite-Alginate-Bone ECM was investigated via SEM imaging with the porosity of the LAP-free (**Figure 2ci-ii**) samples observed to be significantly higher than 3% LAP (**Figure 2ciii-iv**) and 4% LAP (**Figure 2cv-vi**) samples.

Rheological measurements of LAB inks were determined to investigate printing capacity and stability following extrusion. Viscosity was measured as a function of shear rate (**Figure 3a**) confirming a correlation between viscoelastic properties and nanoclay concentration, with higher viscosity at different shear rates compared to controls (**Supplementary Figure 2a**). LAP inclusion augmented viscosity in all blends (**Figure 3a**,**b**) across the range of shear rates examined. The increase in LAP concentration was found to significantly enhance viscous moduli of nanocomposites at a fixed shear rate (**Figure 3c**) confirming the ability of the nanoclay to enhance the viscous properties of the poorly viscous polymers. Storage and Loss moduli of the nanocomposite blends (**Figure 3d**,**i-iv**) displayed a viscoelastic behaviour compared to the controls (**Supplementary Figure 2b**) with stable storage and loss moduli as the angular momentum was increased.

**Figure 3.**
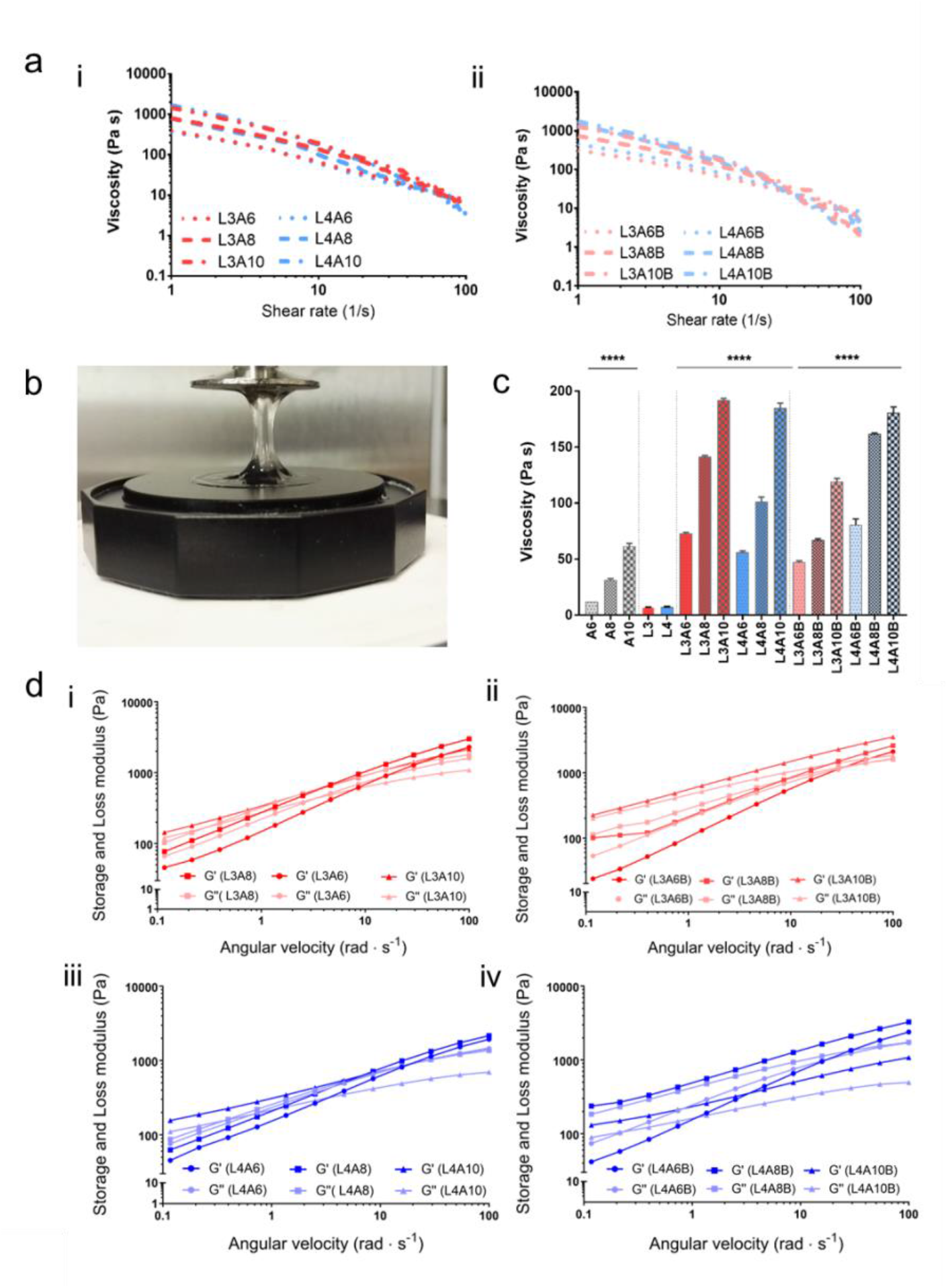
Rheological characterisation of nanoclay-based bone ECM inks. (a) Viscosity over shear rate study of a series of nanoclay-based materials in absence (a-i) or inclusion of bone ECM (a-ii). (b) LAB gel over rheometer plates showing viscoelastic behaviour. (c) Viscosity comparison at fixed shear rate. (d) Storage and loss moduli of nanoclay-based materials without (d-i,iii) and when blended with bone ECM (d-ii,iv). Statistical significance assessed by one-way ANOVA. Mean ± S.D. n=3, ****p<0.00001

### 3.2 Printing characterisation of nanocomposite bone ECM ink

To evaluate the printing resolution and shape fidelity of the nanocomposite bone ECM inks, a regular pattern with increasingly spaced fibre distances was generated. A custom G code was written to investigate the ability of the inks of different LAP and alginate composition to be deposited as fine fibres deposited at an incrementally greater distance of 200 µm steps. The length of the fused portion of printed fibres (fs) and fibre thickness (ft) were measured and the resulting quotients plotted against fibre distance (fd). Micrographs (**Figure 4**) were analysed following AB (**Figure 4a**) and LAB (**Figure 4b**) deposition.

**Figure 4.**
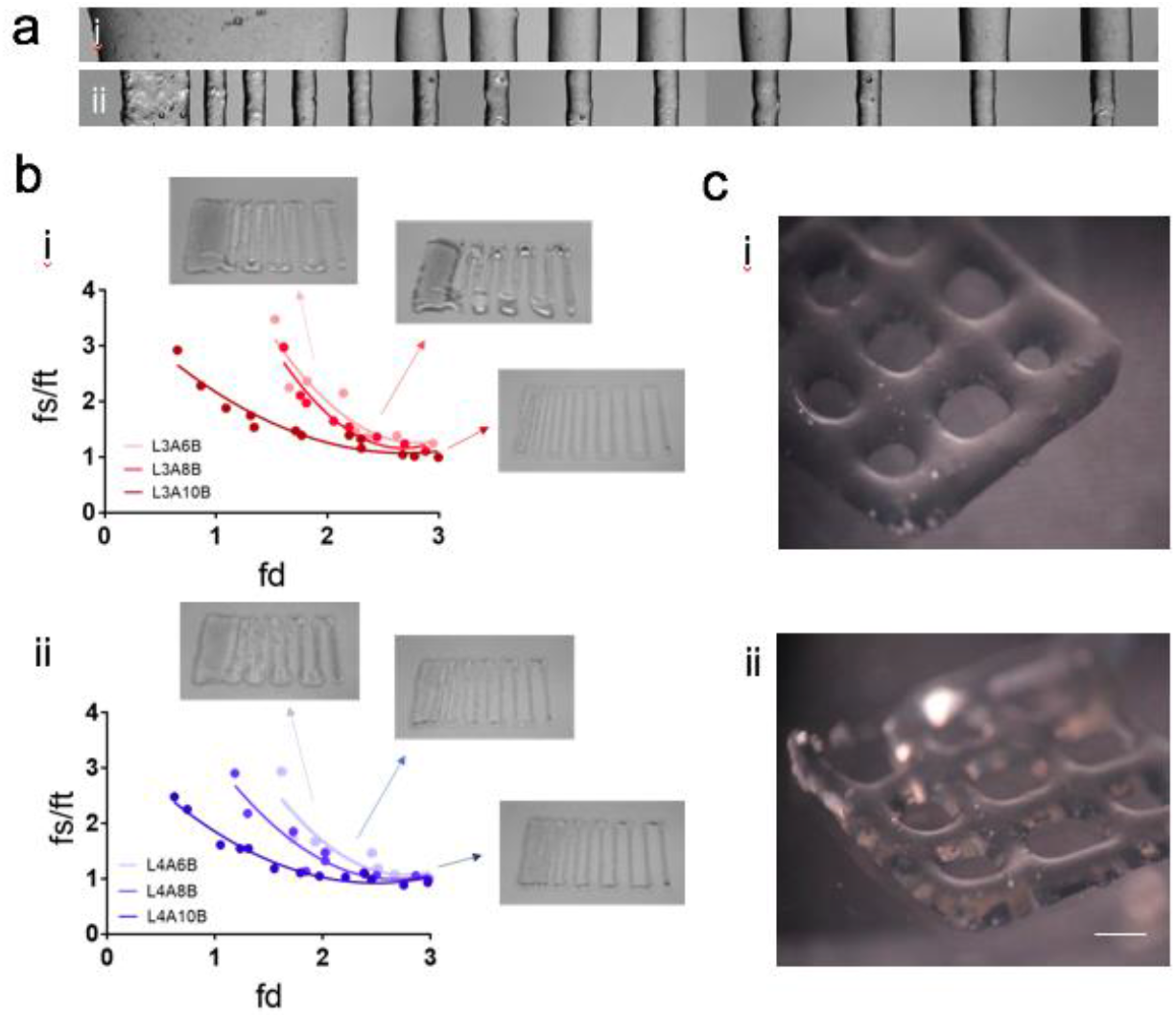
Printing fidelity of nanocomposite bone ECM inks. Filament fusion test (a) was carried out with (a-i) AB and (a-ii) LAB inks. Measurements of the filament fusion tests performed with 3% (b-i) and 4% (b-ii) LAP composite inks. Micrographs of scaffolds printed with 3% (c-i) and 4% (c-ii) LAP-based inks. Scale bar: (c) 1mm.

The resulting analysis indicated that the inclusion of increasingly greater percentage of alginate (6%, 8% and 10% w/v) in inks with fixed Laponite content 3% w/v (**Figure 4b-i**) and 4% w/v (**Figure 4c-i**) augmented the printability of the nanocomposite formulation. The rapid decrease in measured values with incremental change in fibre distance, confirmed enhanced shape fidelity and resolution. The nanocomposite bone ECM ink comprising 3% w/v nanoclay was found to be printable and could be deposited with consistency over 4 layers (**Figure 4b-ii**). The inclusion of an increased percentage of nanoclay (4% w/v) facilitated the printing of increasingly stable scaffolds (**Figure 4c-ii**) at low concentration of alginate. Consequently, a concentration of 4% w/v LAP and 8% w/v alginate was used in all functional studies.

### 3.3 Nanocomposite bone ECM inks support HBMSCs retention, viability and functionality after printing

HBMSCs were encapsulated in nanoclay-free ink as control and printed in nanocomposite bone ECM hydrogel, to investigate the viability of HBMSCs following deposition in 3D. Viability was investigated in control (**Figure 5a**,**b**,**c**,**i-iii**) and nanocomposite LAB ink (**Figure 5d**,**e**,**f**,**i-iii**) using a live/dead assay. Cells remained viable in 3D printed scaffolds (**Figure 5g**) at D1 (83.50 ± 2.23 % and 89.82 ± 3.17 %), D7 (84.78 ± 1.46 % and 90.53 ± 4.50 %) and D21 (80.05 ± 6.67 % and 91.72 ± 3.48 %) in AB and LAB, respectively. Proliferation of printed HBMSCs was subsequently quantified over 21 days of culture *in vitro*. HBMSCs printed in LAP-free ink were observed to proliferate for up to 7 days post-printing comparable to nanocomposite ink samples. After 21 days, HBMSCs density decreased significantly in AB ink, compared to LAB material, which, was found to sustain a low but steady cell growth over 21 days.

**Figure 5.**
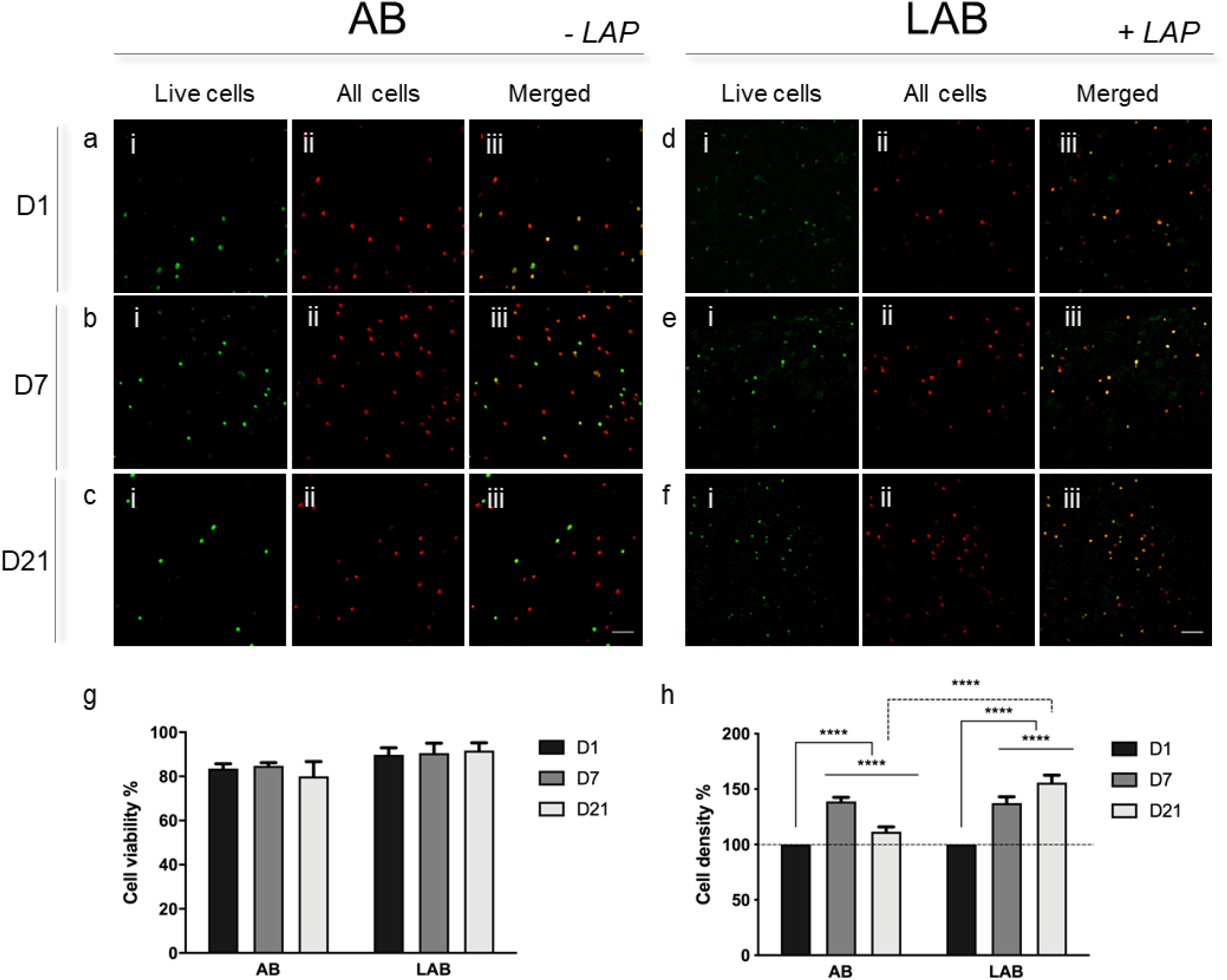
HBMBSCs post-print viability and proliferation. Live-dead assay was performed on 3D printed AB (a-c) and LAB (d-f) scaffolds at 1, 7 and 21 days. Cell viability (g) and density (h) quantification following Image J analysis. Scale bars: (a-f) 100 µm. Statistical significance determined using two-way ANOVA. Mean ± S.D. n=3, ****p<0.00001

To confirm the osteogenic potential of specific nanocomposite blends, ALP staining and analysis was performed on HBMSCs cultured on 2D films of LAP-based bone ECM hydrogels (**Supplementary Figure 3-5**) as well as on cell culture plastic. Culture in basal and osteogenic media revealed an enhanced temporal ALP deposition by HBMSCs on LAB blends with varying concentrations of Laponite (L3, **Supplementary Figure 3**; L4, **Supplementary Figure 4**) over 7 days compared to controls (**Supplementary Figure 5**). LAP materials were observed to support HBMSCs differentiation at early stages (day 1) when seeded at high density.

HBMSC-laden bone ECM inks were 3D printed and cultured for up to 21 days in basal and osteogenic culture media. Printed nanoclay-free bioink (AB), compared to cell-free controls and cell-laden LAB scaffolds, showed limited expression of ALP both at day 1 (**Supplementary Figure 6 a-i, b-i**), 7 (**Supplementary Figure 6 a-ii, b-ii**) and 21 (**Supplementary Figure 6 a-iii, b-iii**) in basal and osteogenic conditions. The inclusion of LAP within the material ink was found to elicit an enhanced deposition of ALP at day 1 (**Supplementary Figure 6 c-i, d-i**), day 7 (**Supplementary Figure 6 c-ii, d-ii**) and 21 (**Supplementary Figure 6 c-iii, d-iii**) in basal and osteogenic media.

### 3.4 The inclusion of nanoclay in bone ECM inks improved drug retention and sustained release

To evaluate the ability of nanoclay bone ECM inks to retain biologics/compounds of interests such as lysozyme, bovine serum albumin, BMP-2 and VEGF, the agents were adsorbed onto 3D printed scaffolds for 24h. Following adsorption, *in vivo* conditioning was simulated by adding a collagenase solution able to trigger material degradation, therefore forcing the release of the absorbed cargo.

The ability of LAB and LAP-free scaffolds (AB) to absorb and retain biologics of interests, was examined by the quantification of kinetic release of lysozyme (**Supplementary Figure 7a**) and BSA (**Supplementary Figure 7b**) over 48h. LAB adsorbed a greater concentration of both lysozyme and BSA. Collagenase inclusion after 24h adsorption, triggered the release of the cargo agents, enabling LAP-based scaffolds to retain a significantly larger proportion of lysozyme and BSA compared to AB for up to 24h

To investigate the ability of the 3D printed LAB scaffold to retain and localise growth factors of interest for bone regeneration, VEGF was adsorbed by 3D printed LAB and AB controls and implanted in the developing chick embryo CAM (**Figure 6a-i**,**v**). The explanted groups were observed to be highly vascularised (**Figure 6b-i**,**v**), evidenced by Chalkley score analysis (**Figure 6c**). LAB scaffolds loaded with VEGF contained a significantly greater number of vessels (p<0.0001) compared to empty, AB-VEGF and VEGF-free controls (AB and LAB) implanted scaffolds. Histological analysis (**Figure 6d-g**) confirmed the potential of VEGF-loaded samples to promote blood vessel formation as well as a higher deposition of collagenous matrix in LAP-based VEGF-loaded groups.

**Figure 6.**
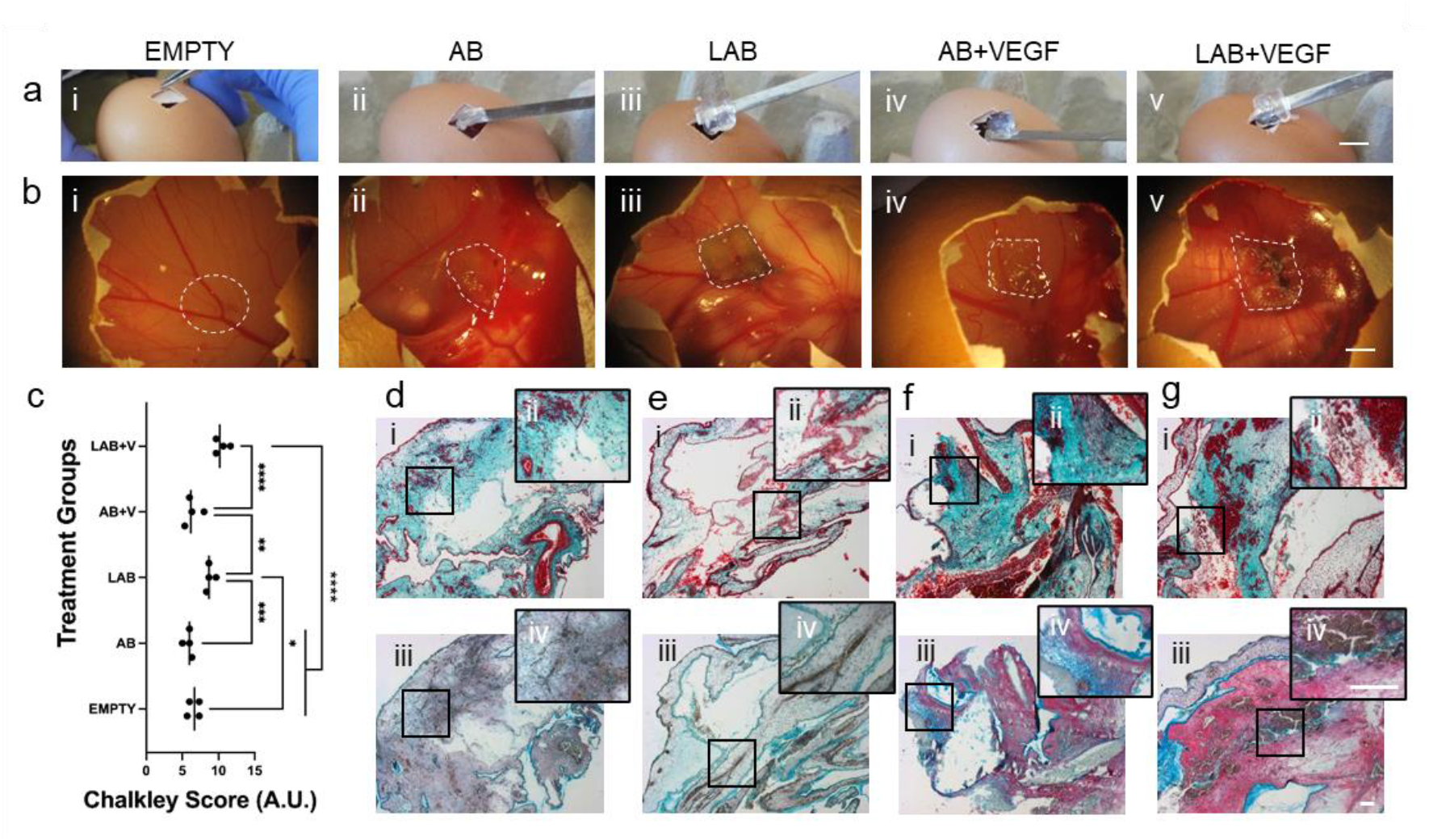
Nanoclay-based ink support sustained release of VEGF in CAM model. (a) Macrographs during sample implantation and (b) retrieval. (i) Empty, (ii) AB, (iii) LAB, and VEGF-loaded (iv) AB and (v) LAB 3D printed scaffolds. (c) Chalkley score of vascularised samples and controls. (d) Histological micrographs of samples stained for (i-ii) Goldner’s Trichrome and (iii-iv) A&S. Statistical significance assessed by one-way ANOVA. Mean ± S.D. n=4, *p<0.05, **p<0.01, ***p<0.001, ****p<0.0001. Scale bar: (a,b) 10mm, (d-g) 100µm.

Additional CAM analysis was undertaken to explore the synergistic effect of HBMSCs and BMP-2 in an *ex vivo* scenario. Compared to empty controls (**Figure 7a**) the implanted 3D printed constructs LAP-free (**Figure 7b-i**,**iv**) and the nanoclay-based (**Figure 7c-i**,**iv**) constructs were observed to be fully integrated. Blood vessels were quantified using the Chalkley score method (**Figure 7d**). HBMSC-laden LAB scaffolds containing BMP-2 were highly vascularised with increased blood vessels present compared to HBMSC-laden BMP-2-loaded AB scaffolds (p<0.001), empty controls and LAP-free acellular and BMP-2-free scaffolds (p<0.0001). LAB scaffolds were found to promote significant vascularisation compared to AB scaffolds (p<0.01).

**Figure 7.**
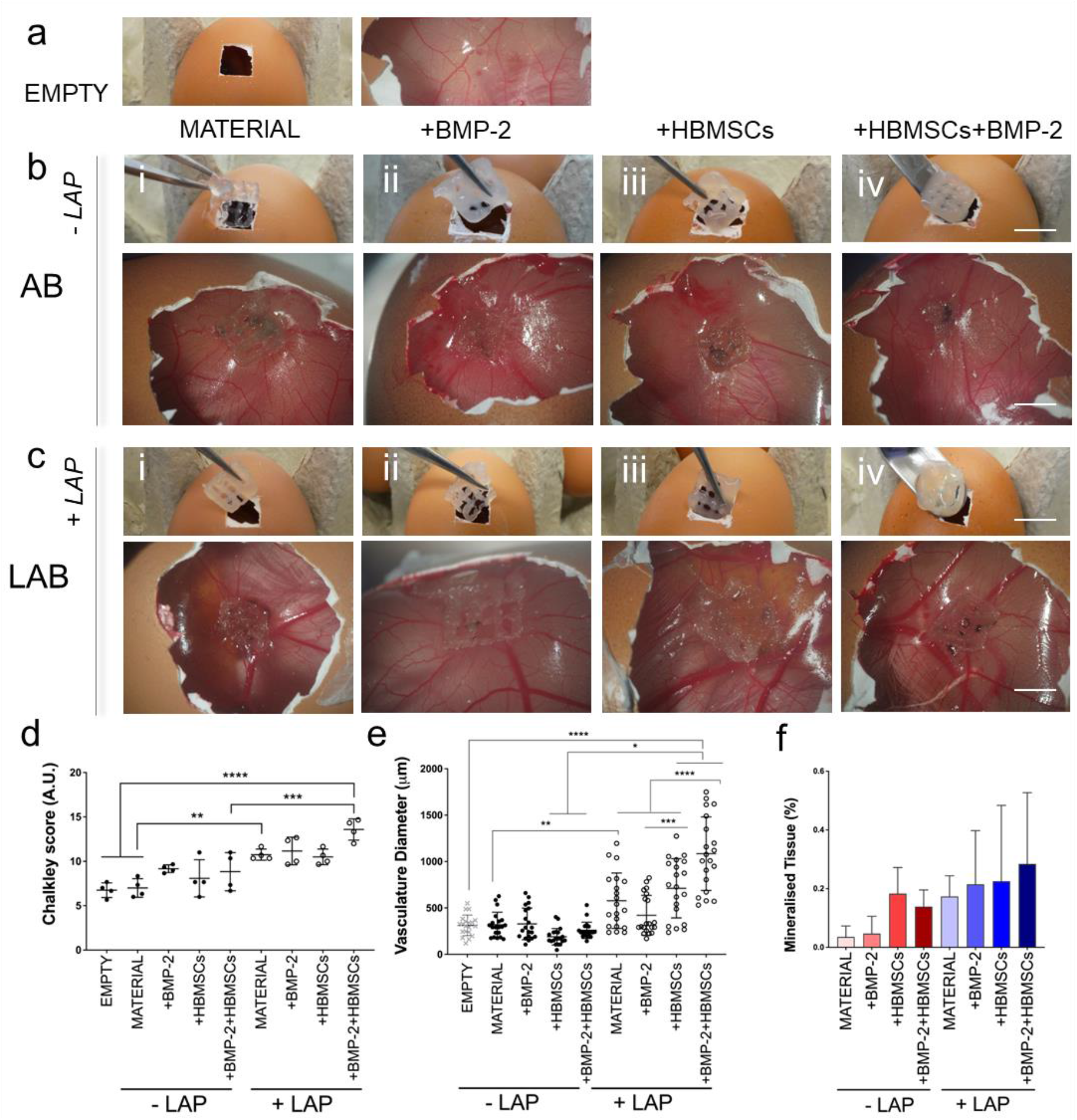
Nanocomposite bone ECM scaffolds support mineralisation *ex vivo*. (a) Macro- and micro-graphs of empty control. Implanted and explanted (b) LAP-free and (c) LAP-loaded 3D (i) material (drug- and cell-free) control, (ii) BMP-2 loaded, (iii) cell-loaded and (iv) BMP-2 and cell loaded scaffolds. (d) Chalkley score of implanted samples and control after 7 days of culture. (e) Quantitative analysis of afferent vascular supply to implanted scaffolds before extraction. (f) u-CT analysis of implanted scaffolds following 7 days of incubation in a CAM model. Scale bar: (a,b,c) 10mm Statistical significance assessed by one-way ANOVA. Mean ± S.D. n=4, *p<0.05, ** p<0.01, ***p<0.001, ****p<0.0001

Vessel diameters were measured *in ovo* prior to isolation. LAB acellular and biologic scaffolds were observed to be significantly larger (p<0.01) than AB 3D printed materials. The inclusion of LAP nanosilicate discs significantly enhanced blood vessel diameter (p<0.0001) when combined with BMP-2, HBMSCs and both. Thus, the synergistic combination of HBMSCs and BMP-2 was found to stimulate the formation of larger vessels (1 mm) compared to control AB and LAB scaffolds (p<0.0001). Micro-CT analysis of explanted 3D scaffolds revealed the presence of mineralised tissue although this was not significantly greater than control acellular and BMP-2 free printed inks.

Histological analysis (**Figure 8**) revealed vascularisation of samples in LAP-free (**Figure 8a-d**) and LAP-based constructs (**Figure 8e-h**). Implanted nanoclay-free 3D constructs loaded with BMP-2 and HBMSCs (**Figure 8d-i**,**ii**) resulted in leakage of vessels in the chorioallantoic membrane, resulting in an extensive penetration of vessels accompanied by erythrocytes dispersion across the implant. A collagenous matrix was present in cell-laden groups (both LAP-free and LAP-based), demonstrating the functionality of HBMSCs after 7 days of implantation. LAP-based controls stained positive for the mineral stain von Kossa compared to LAP-free controls.

**Figure 8.**
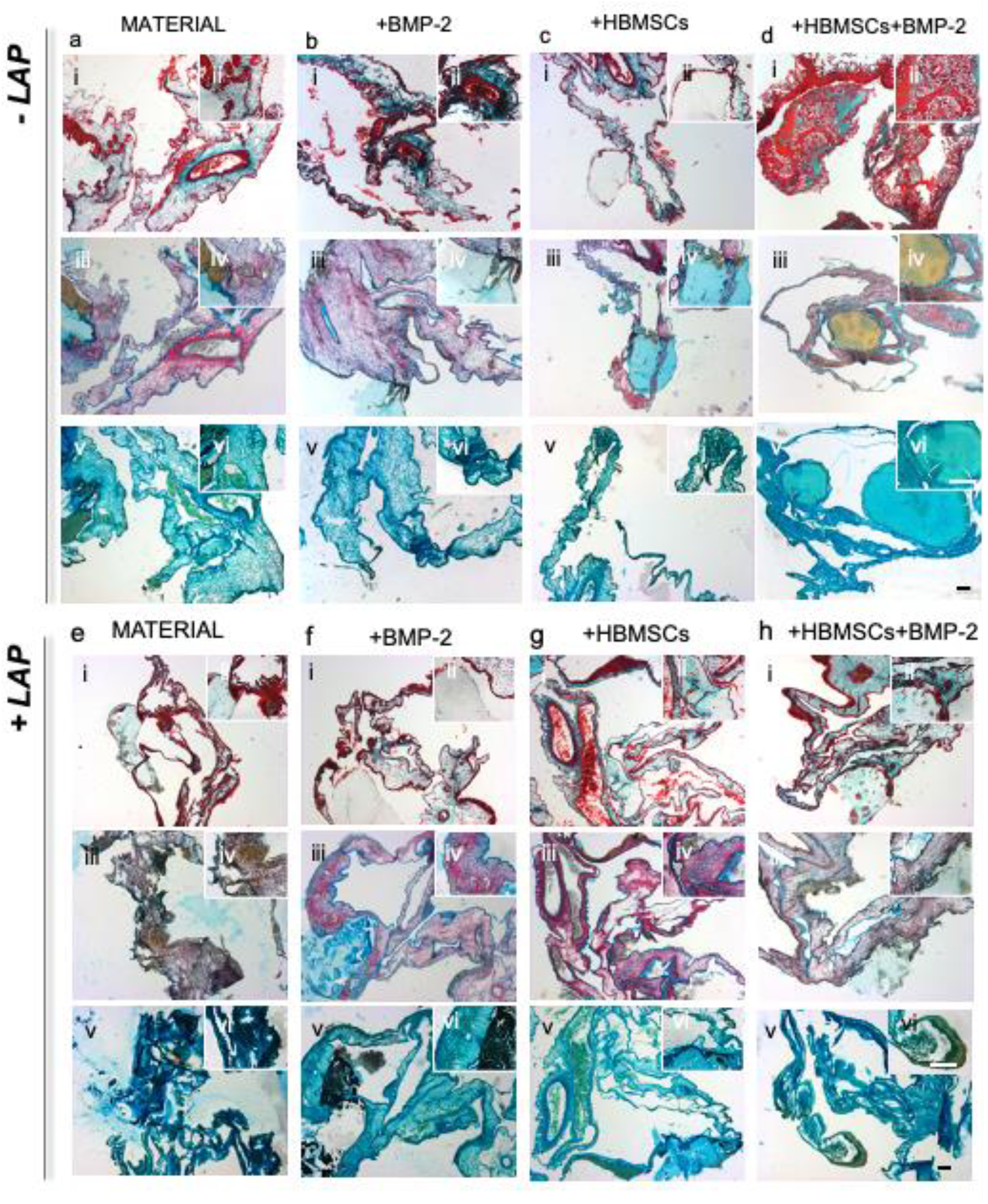
CAM implantation of 3D printed scaffolds containing BMP-2 and HBMSCs. LAP-free (a-d) and LAP-loaded (e-h) groups are stained for Goldner’s Trichrome (i,ii), Alcian Blue & Sirius Red (A&S) (iii,iv) and Von Kossa (v, vi). Scale bars: 100µm.

## 4. Discussion

The use of biofabrication technologies has advanced the engineering of 3D substitutes for the repair of damaged and diseased skeletal tissue. However, the lack of functional inks, capable of supporting cell growth and differentiation post-printing and, ultimately, to regenerate skeletal defects, remains an unresolved challenge. The current study details the incorporation of human demineralised and decellularized bone ECM in combination with nanosilicate (Laponite^®^, LAP) particles and alginate polymer for the design of a bioactive ink. The addition of both LAP and alginate to a human bone decellularised and demineralised ECM was found to stabilise the sol fraction and the mass swelling ratio at low polymeric content.

The engineering of nanocomposite materials, incorporating functional fillers capable of modifying physical properties (e.g. thixotropic behaviour), compound interactions (e.g. drug localisation) and biological functionality (e.g. cell spreading) has supported the fabrication of cell-laden constructs for the active repair of skeletal defects. Nevertheless, the sole inclusion of nano-fillers does not guarantee the engineering of a functional microenvironment for stem/progenitor cell proliferation and, ultimately, differentiation ^27^. Decellularised ECM provide a particularly attractive approach to mimic the native tissue-specific micro-environment. Recently, a number of studies ^17,28,29^ have demonstrated the ability to print non-human decellularised ECM (particularly cardiac ^17,28^ and hepatic ^29^ tissues) in combination with clay nanodiscs, demonstrating the beneficial inclusion of nanoclay fillers to drastically improve printability and print fidelity. Nevertheless, the animal-sourced decellularised materials (mainly porcine) while providing a similar collagen, glycosaminoglycans and growth factors content, can still generate an immune response. Thus, human-based decellularised tissue has come to the fore as an ideal biomaterial for tissue regeneration.

The investigation of the microstructure of the LAB material revealed a difference in porosity. LAP-based inks were found to be less porous as the positive rim charge of the nanoparticles can closely interact with negatively charged alginate and collagen-abundant bone ECM components. This was further confirmed by rheological studies, demonstrating a significant increase in viscous properties with the inclusion of nanoclay particles within the composites behaviour already observed in a number of previous studies ^10,12,27^. Indeed, LAP nanoparticles hold the ability to closely interact electrostatically with polymeric chains, reducing the distance between the biomaterial network thus increasing viscosity and ultimate mechanical properties. Moreover, the shear-thinning properties of LAP-based inks have been found essential for 3D bioprinting applications of skeletal implants ^12^. The control over viscoelastic properties and the influence on printability, was demonstrated by the filament fusion test. Results highlighted the ability of an increased concentration of LAP to significantly influence printability over a number of stacked layers. However, alternate 0°/90° patterning was observed to be influenced by the post-printing relaxation of the viscous properties, with an increase in shape fidelity directly correlated with increase of LAP content in agreement with was previously reported ^9,30^.

The overall viscoelastic properties of the LAB ink were tuned to allow the printing of HBMSCs. LAP-based cell-laden scaffolds supported HBMSCs proliferation over 21 days compared to LAP-free control as previously reported ^10,12^. The cell retention ability of LAB scaffolds was a likely result of the enhanced viscoelastic properties compared to AB constructs, preserving the integrity of the overall printed construct over time and avoiding the release of cell material from the degrading fibres. Furthermore, in agreement with previous results ^9,11^, LAP inclusion was found to aid HBMSCs differentiation towards bone lineage, as highlighted by the ALP staining micrographs. The 3D printing of HBMSCs reduced spatial spreading of encapsulated stromal cells and facilitated a functional response with intense ALP expression *in vitro* as well as collagen deposition following *ex vivo* implantation, as previously reported^10,11^.

Indeed, the addition of LAP nanodiscs facilitated the local release of ALP over 21 days. As previously reported ^11,12^, nanocomposite inks stimulated ALP deposition immediately after printing (day 1), supporting the rapid formation of skeletal-specific scaffolds. Thus, the nanosilicate inclusion detailed in these studies offers an approach for *in vitro* bone modelling with the combination of alginate, specifically supporting the 3D deposition, while addition of bone ECM enhanced the functionality of the printed scaffold. ALP was found to be expressed ubiquitously in nanoclay-based sample groups as previously reported^10–12^. The deposition of ALP was correlated with the concentration of LAP in the composite (Supplementary Figure 3 and 4). The ALP staining in LAP-only and LAP-bone ECM samples was present from day 1 to day 7 both in basal and osteogenic conditions. Nevertheless, the presence of alginate appeared to alter the morphology of seeded HBMSCs as previously reported ^9,27^ with HBMSCs developing a rounded-morphology and concomitant expression of ALP observed from the first day of culture.

The ability of nanoclay-modified bone-ECM scaffolds to localise biological agents of interest within a preclinical scenario was investigated using the CAM assay. *Ex vivo* implantation of LAP-based bone ECM 3D printed constructs demonstrated the ability of the new blend to support angiogenesis, with vessels forming over 7 days of implantation, as a consequence of the localisation of GFs within the matrix. The retention of VEGF was found to stimulate vessel ingrowth in LAP-based implants, as previously demonstrated for constructs comprising the nanoclay material ^10^. Furthermore, the current studies illustrate the synergistic interaction of a nanocomposite ink (LAP and alginate) microenvironment for HBMSCs proliferation and functionality. Indeed, the deposition of cell-laden BMP-2-loaded constructs resulted in enhanced mineralisation and vascularisation. In addition, the diameter of the CAM blood vessels was significantly increased when LAP was combined with alginate and bone ECM. This biological observation is well documented for local BMP-2 exposure ^12^, but less clear in drug-free implants. Thus, bone ECM in combination with LAP was found to support angiogenesis, providing a platform to stimulate vascularisation of a skeletal TE construct. On-going work in our laboratories will address the underlying biochemical mechanisms.

We are cognisant that the use of human bone ECM tissue could be initially limited by legal and ethical restrictions impacting on clinical translation. However, the possibility to generate patient-specific decellularised bone ink, harnessing the patients’ own skeletal tissue, is appealing and offers an exciting opportunity for a personalised medicine approach to aid bone repair harnessing a native engineered tissue substitute.

## 5. Conclusions

The design of biomimetic functional biomaterials for skeletal tissue engineering is a key goal to aid bone repair. The use of xenogeneic ECM matrices that incorporate GFs and native polymers offers an approach to stimulate an effective repair of the damaged skeletal tissue. However issues around immunogenicity, synthesis and limited mechanical properties have limited the use of ECM matrices for 3D bioprinting purposes.

This study sought to harness human bone-ECM in combination with alginate and nanoclay particles, to fabricate implantable constructs, capable of supporting and promoting bone repair. Results show that LAP limited the swelling of printable inks, enabled tuning of rheological properties and allowed the printing of self-sustained 3D structures, comprising bone-ECM, with an ultra-low polymeric concentration. The novel human bone-ECM ink generated supported the deposition of HBMSCs, maintaining viability and supporting proliferation as well as differentiation along the osteogenic lineage *in vitro and ex vivo*. LAP-based scaffolds were found to retain VEGF or BMP-2 in an *ex vivo* CAM model, highlighting the ability to sustain angiogenic and osteogenic development important in endochondral ossification and skeletal repair. Future studies outside the scope of the current work, will be examine the *in vivo* application of the ECM-based 3D bioprinted skeletal construct, targeting the functional repair of fracture and calvarial preclinical models of bone repair.

In summary, the current study demonstrated the patterning in 3D of a novel nanocomposite ink containing human bone ECM components, capable of supporting cell viability and sustaining growth factor release with potential application for the repair of bone defects.

## Credit author statement

**Yang-Hee Kim:** Methodology, Validation, Investigation, Writing original draft, Funding acquisition. **Janos M Kanczler:** Methodology, Investigation, Writing - review & editing. **Stuart Lanham:** Methodology, Investigation, Writing - review & editing. **Andrew Rawlings:** Methodology, Investigation. **Marta Roldo:** Writing - review & editing. **Gianluca Tozzi:** Writing-review & editing. **Jonathan I. Dawson:** Supervision, Writing - review & editing, Funding acquisition. **Gianluca Cidonio:** Conceptualization, Methodology, Validation, Writing original draft, review & editing, Funding acquisition. **Richard O.C Oreffo:** Conceptualization, Methodology, Supervision, Writing review & editing, Funding acquisition.

## Acknowledgements

This study was supported by grants from the Biotechnology and Biological Sciences Research Council (BBSRC LO21071/ and BB/L00609X/1) and UK Regenerative Medicine Platform Hub Acellular Approaches for Therapeutic Delivery (MR/K026682/1) Acellular Hub, SMART materials 3D architecture (MR/R015651/1) and the UK Regenerative Medicine Platform (MR/L012626/1 Southampton Imaging) to RO and MRC-AMED Regenerative Medicine and Stem Cell Research Initiative (MR/ V00543X/1) to JID and YK. GC acknowledges funding from AIRC Aldi Fellowship under grant agreement No. 25412.

## Data availability

Data will be made available on request.

