## Supplementary material for "Biofabrication of nanocomposite-based scaffolds containing human bone extracellular matrix for the differentiation of skeletal stem and progenitor cells": LAB_3DP_v07-04-2023_Supplementary.pdf

### Supplementary Information

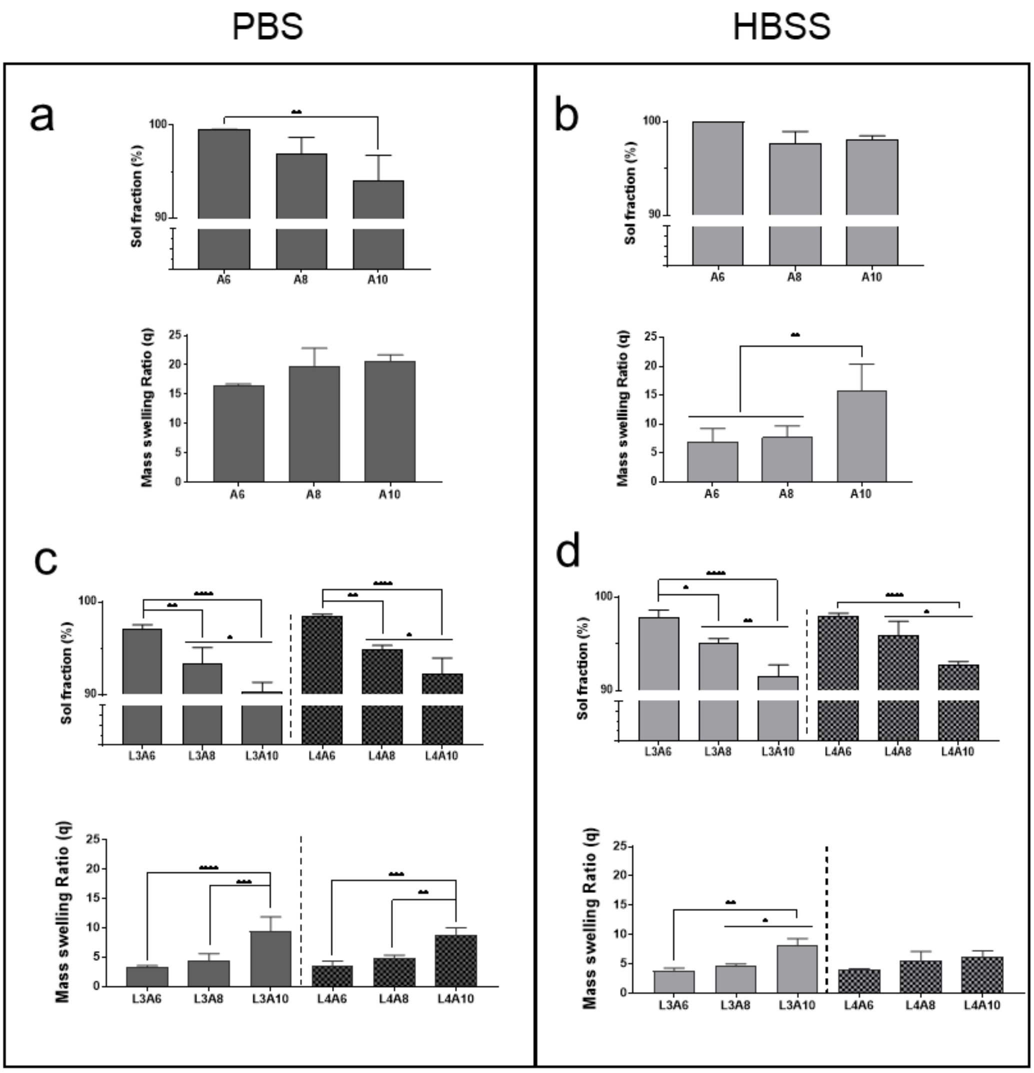


**Supplementary Figure 1.** Physical investigation of control composite inks. Sol fraction and mass swelling ration of alginate and nanoclay-modified alginate controls in PBS (a,c) and HBSS (b,d), respectively. Statistical significance assessed by one-way ANOVA. Mean ± S.D. n=3, ∗p<0.05, ∗∗p<0.01, ∗∗∗p<0.001, ∗∗∗∗p<0.00001

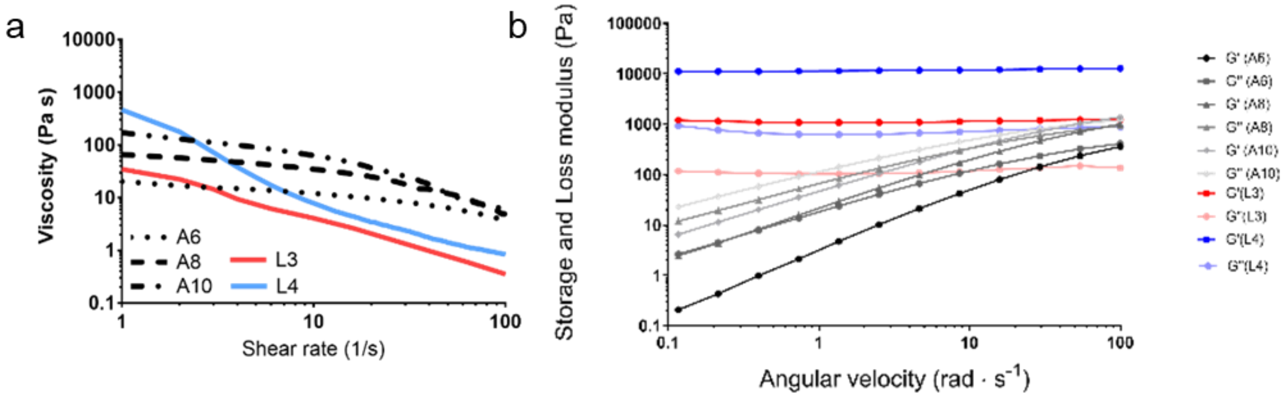


**Supplementary Figure 2.** Rheological characterisation of control materials. Viscosity (a) and storage and loss moduli (b) of Alginate and Laponite controls.

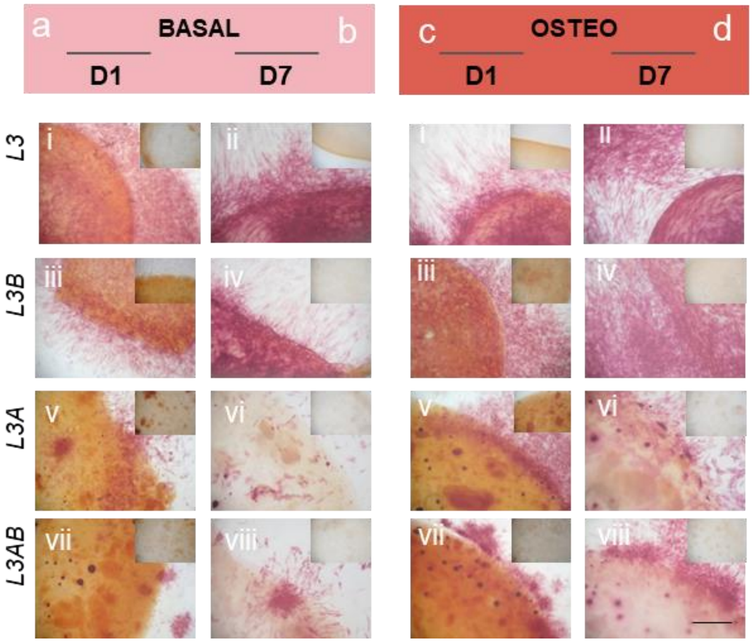


**Supplementary Figure 3.** ALP staining of HBMSCs seeded on 3% w/v nanoclay (Laponite) and composites (Laponite-bone-ECM (L3B), Laponite-alginate (L3A), Laponite-alginate-bone-ECM (L3AB)) at day 1 and 7 both cultured in basal (a,b) and osteogenic (c,d), respectively. Scale bars: 250 µm

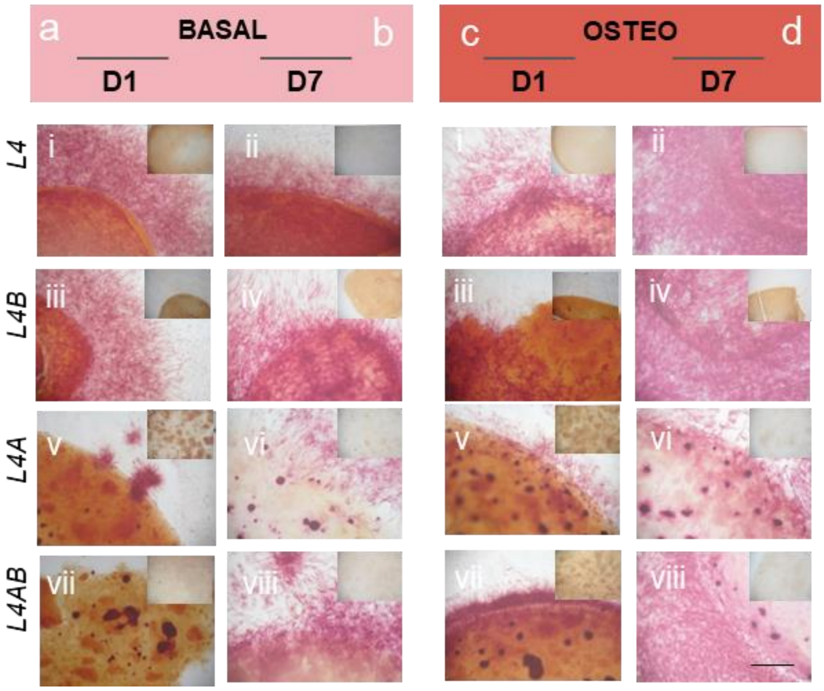


**Supplementary Figure 4.** ALP staining of HBMSCs seeded on 4% w/v nanoclay (Laponite) and composites (Laponite-bone-ECM (L4B), Laponite-alginate (L4A), Laponite-alginate-bone-ECM (L4AB)) at day 1 and 7 both cultured in basal (a,b) and osteogenic (c,d), respectively. Scale bars: 250 µm

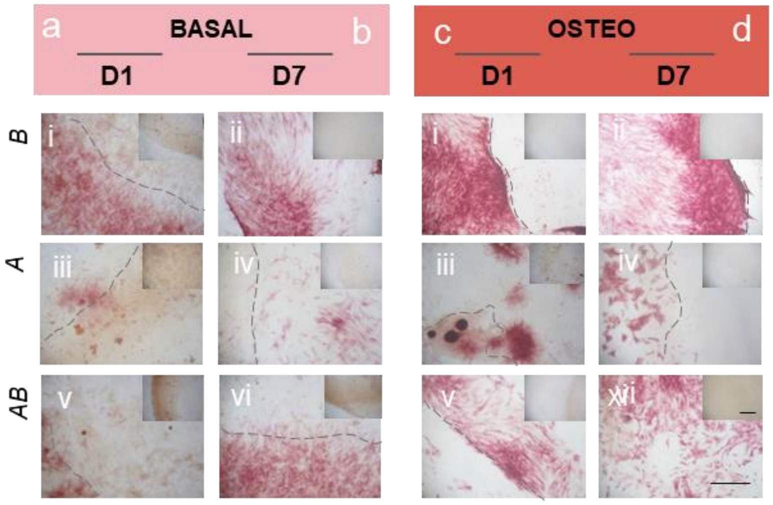


**Supplementary Figure 5.** ALP staining of HBMSCs seeded on bone ECM (B), alginate (A) and composite (AB) at day 1 and 7 both cultured in basal (a,b) and osteogenic (c,d), respectively. Scale bars: 250 µm

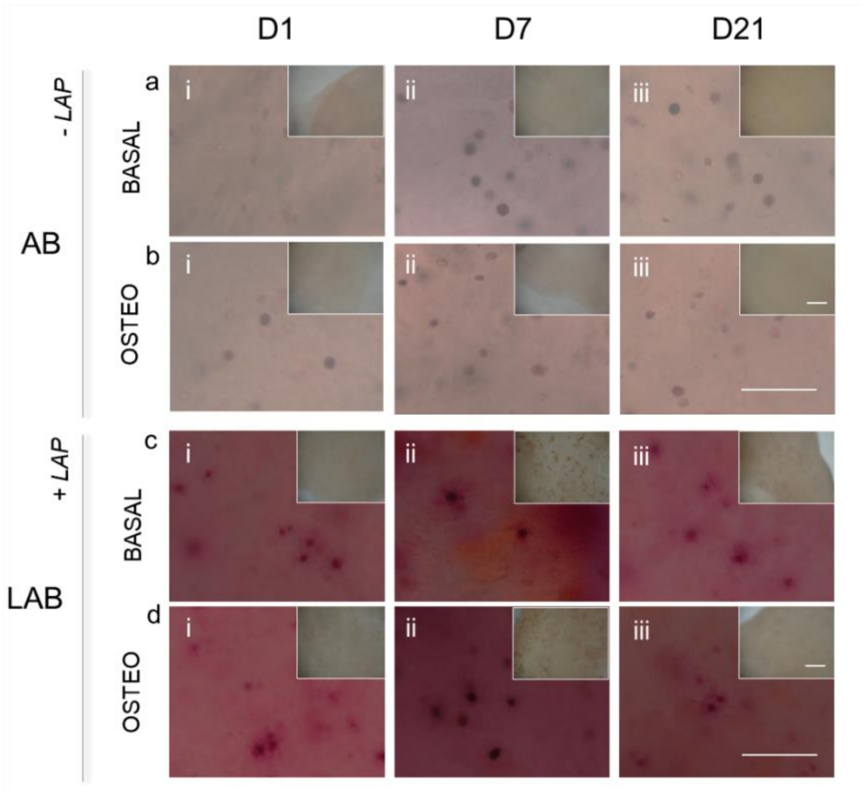


**Supplementary Figure 6.** HBMBSCs printing functionality. Alkaline phosphatase staining of 3D bioprinted scaffolds following cultivation in basal (AB, a; LAB, b) and osteogenic (AB, b; LAB, d) media conditioning complete with acellular control. Scale bars: 50 µm (samples), 250 µm (acellular controls)

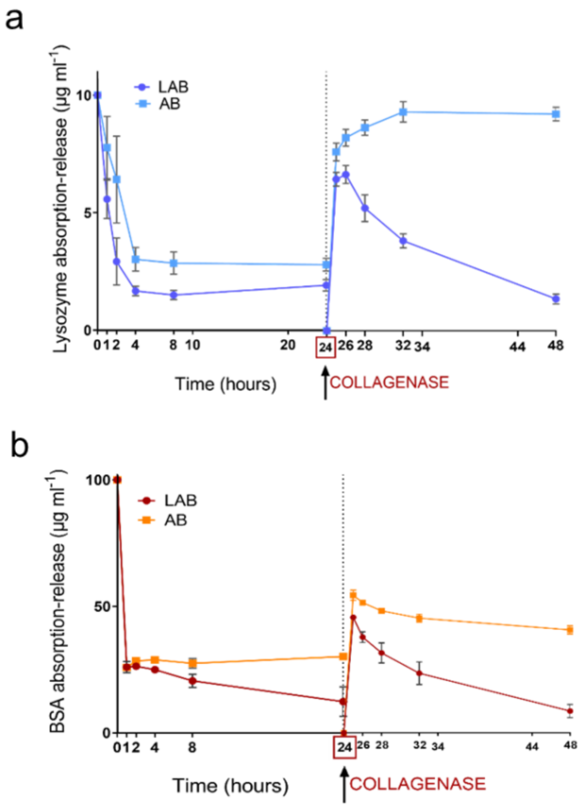


**Supplementary Figure 7.** Absorption/release of model proteins. Lysozyme (a) and bovine serum albumin (b) collagenase-mediated release following 24h absorption.
